# Synergetic effect of non-complementary 5’ AT-rich sequences on the development of a multiplex TaqMan real-time PCR for specific and robust detection of *Clavibacter michiganensis* and *C. michiganensis* subsp. *nebraskensis*

**DOI:** 10.1101/566281

**Authors:** Adriana Larrea-Sarmiento, Anne M. Alvarez, James P. Stack, Mohammad Arif

## Abstract

*Clavibacter* is an agriculturally important genus comprising a single species, *Clavibacter michiganensis*, and multiple subspecies, including, *C. michiganensis* subsp. *nebraskensis* which causes Goss’s wilt/blight of corn and accounts for high yield losses - listed among the five most significant diseases of corn in the United States of America. Our research objective was to develop a robust and rapid multiplex TaqMan real-time PCR (qPCR) to detect *C. michiganensis* in general and *C. michiganensis* subsp. *nebraskensis* with enhanced reliability and accuracy by adding non-complementary AT sequences to the 5’ end of the forward and reverse primers. Comparative genomic analyses were performed to identify unique and conserved gene regions for primer and probe design. The unique genomic regions, ABC transporter ATP-binding protein CDS/ABC-transporter permease and MFS transporter were determined for specific detection of *C. michiganensis* and *C. m.* subsp. *nebraskensis*, respectively. The AT-rich sequences at the 5’ position of the primers enhanced the reaction efficiency and sensitivity of rapid qPCR cycling; the reliability, accuracy and high efficiency of the developed assay was confirmed after testing with 59 strains from inclusivity and exclusivity panels – no false positives or false negatives were detected. The assays were also validated through naturally and artificially infected corn plant samples; all samples were detected for *C. michiganensis* and *C. m*. subsp. *nebraskensis* with 100% accuracy. The assay with 5’ AT-rich sequences detected up to 10- and 100-fg of *C. michiganensis* and *C. michiganensis* subsp. *nebraskensis* genome targets, respectively. No adverse effect was observed when sensitivity assays were spiked with host genomic DNA. Addition of 5’ AT rich sequences enhanced the qPCR reaction efficiency from 0.82 (M = -3.83) and 0.91 (M = -3.54) to 1.04 (with optimum slope value; M = -3.23) for both *C. michiganensis* and *C. michiganensis* subsp. *nebraskensis*, respectively; a increase of 10-fold sensitivity was also obtained with *C. michiganensis* primer set. The methodology proposed here can be used to optimize the reaction efficiency and to harmonize the diagnostic protocols which have prodigious applications in routine diagnostics, biosecurity and microbial forensics.

## Introduction

The gram-positive aerobic species *Clavibacter michiganensis*, which resides in the xylem vessels of the host, is a devastating bacterial pathogen of many economically important agricultural crops. Currently, the genus *Clavibacter* is known only for one species, *C. michiganensis* which belongs to family *Microbacteriaceae* within the phytopathogenic coryneform group, known for epi- or endophytic colonization of symptomatic as well as several asymptomatic plant species [1,2]. Virulence genes within this species are plasmid-borne (not essentially present), while host colonizing genes are chromosomal [3]. Hitherto, nine subspecies based on host specificity and bacteriological characteristics have been identified within the single species of *C. michiganensis.* Four subspecies infect two important plant species in the *Solanaceae* family (tomato and pepper) while the other five subspecies infect plant hosts from *Fabaceae, Solanaceae* and *Poaceae* families - *C. m.* subsp. *michiganensis* (*Cmm*; bacterial canker of tomato), *C. m.* subsp. *capsici* (*Cmc;* bacterial canker of pepper), *C. m.* subsp. *insidiosus* (*Cmi;* stunt and wilt of alfalfa), *C. m.* subsp. *nebraskensis* (*Cmn;* wilt and blight of maize), *C. m.* subsp. *sepedonicus* (*Cms;* ring rot of potato), *C. m.* subsp. *tessellarius* (*Cmt;* leaf freckles and leaf spots of wheat), *C. m.* subsp. *phaseoli* (*Cmp*; bacterial leaf yellowing of bean), *C. m.* subsp. *californiensis* (*Cmcf*; bacterial infection of tomato seeds) *C. m.* subsp. *chiloensis* (*Cmcl*; bacterial infection of pepper seeds) [1,2,4,5].

*C. michiganensis* subsp. *nebraskensis* is the causal agent of Goss’s bacterial wilt and blight of corn (*Zea mays*), which is listed among the five most significant diseases of corn in the U.S. Goss’s wilt accounts for high yield losses in the Great Plains and Midwest regions of the corn belt of the U.S., the largest producer and exporter of corn worldwide. Since the first report in 1969 from Nebraska, the disease has been spreading and adversely impacting corn production in the Midwestern United States including western Indiana, Illinois, Iowa, Missouri, eastern Nebraska, and eastern Kansas, as well as Canada. Goss’s disease shows two major types/phases of symptoms - a leaf blight and a systemic vascular wilt [6,7,8]. The leaf blight phase involves leaves and above-ground parts of the plant which exhibit small, dark and discontinuous water-soaked spots; orange shiny bacterial exudates are also observed in advanced stages. The less common wilt phase involves systemic infection of xylem elements resulting in discoloration of vascular bundles, stalk rot and premature plant death. Transmission of Goss’s disease usually occurs through open wounds on leaves, and C. *m.* subsp. *nebraskensis* can overwinter and remain in plant debris to serve an inoculum for the next crop; transmission through seeds is minimal [2,8,9].

Thus far, no foliar bactericide applications are available for control; therefore, disease resistance/tolerance genotype selection with integrated pest management practices including crop rotation, deep tillage, and improved drainage are the most effective measures. Diagnosis of Goss’s wilt is limited to recognition of variable symptoms followed by isolation onto semi-selective media and identification of *C. michiganensis* of the subspecies that causes Goss’s wilt using additional tests. Recently, Dobhal and collaborators [10] developed a loop-mediated isothermal amplification (LAMP) method to discriminate all known subspecies of *C. michiganensis*; however, no effective qPCR/PCR protocol is available. Although, serological and molecular diagnostic methods for economically important subspecies of *C. michiganensis* have been developed [2,7,11,12,13,14], these diagnostic tests are time consuming and may exhibit cross reactivity with Gram-positive bacteria and/or Gram-negative bacteria demanding further molecular test for cross confirmation [11,15].

Real-Time qPCR is a powerful and robust technique and offers greater sensitivity and speed compared with conventional PCR; however, qPCR sensitivity can be compromised as a result of poor primer thermodynamics. Primer thermodynamics can be improved by adding 5’ AT-rich sequences often called overhang, flap, or tail [16,17,18]). In addition, primer Tm and length can also be fine-tuned to harmonize the amplification reaction conditions [16,17]. The tailed primers have been reported to provide higher PCR yields and sensitivity [16,17,18]. Unfortunately, the underlying mechanism behind the action of flap primers is still unknown.

In this study, we conducted detailed *in vitro* and *in silico* analyses designed to enhance the effect of tailed primers in TaqMan real-time qPCR assays that simultaneously detect all nine subspecies of *C. michiganensis* as well as specifically identifying *C. m.* subsp. *nebraskensis.*

## Materials and Methods

### Source of Bacterial Strains and DNA Isolation

Thirty-two bacterial strains within the species *C. michiganensis* (including, fourteen *C. m.* subsp. *nebraskensis*, nine *C. m.* subsp. *michiganensis*, two *C. m.* subsp. *sepedonicus*, one *C. m.* subsp. *chiloensis*, two *C. m.* subsp. *capsici*, one *C. m.* subsp. *californiensis. m.* subsp. *californiensis*, one *C. m.* subsp. *phaseoli*, one *C. m.* subsp. *insidiosus* and one *C. m.* subsp. *tessellarius*) and twenty-five phylogenetically related strains including type strains and corn saprophytes, were used in this study (Table 1). Strain designations beginning with “A” were obtained from the Pacific Bacterial Collection at the University of Hawaii at Manoa. Bacterial cultures were taken from - 80°C; gram-positive bacteria were grown on TZC-S medium (peptone 5 g/L, 10 g/L, sucrose 5 g/L and agar at 17 g/L, 0.001% 2,3,5-triphenyl-tetrazolium chloride) and gram-negative bacteria were grown on TZC medium with 5 g/L dextrose as the carbon source and 0.001% tetrazolium chloride) [19]- plates were incubated at 26°C (±2°C). Grown cultures were used for DNA isolation using Wizard Genomic DNA Purification Kit following the manufacturer’s instructional manual (Promega, Madison, WI).

**Table 1.** Fifty-nine strains of plant-associated bacteria were used in inclusivity and exclusivity panels to validate the specificity of multiplex TaqMan real-time PCR developed to detect all *Clavibacter michiganensis* subspecies and specifically, *C. michiganensis* subsp. *nebraskensis*.

| Sample code | Other ID<sup>a</sup> | Accession number | Strain | Host | Origin | Average C<sub>T</sub> value<sup>b</sup> |  |
| --- | --- | --- | --- | --- | --- | --- | --- |
|  |  |  |  |  |  | CM | Cmn11 |
| A6203<sup>c,d</sup> | DP104A | MH560461 | *C. michiganensis* subsp. *nebraskensis* (Cmn) | corn | OK, USA | 19.2 | 23.7 |
| A6204<sup>c,d</sup> | DP114A | MH560462 | *Cmn* | corn | TX, USA | 14.65 | 19.26 |
| A6205<sup>c,d</sup> | DP115 | MH560463 | *Cmn* | corn | IA, USA | 12.03 | 17.68 |
| A6206<sup>a,c</sup> | DP117 | MH560464 | *Cmn* | corn | NE, USA | 11.88 | 17.32 |
| A6207<sup>c,d</sup> | DP121 | MH560465 | *Cmn* | corn | TX, USA | 11.05 | 16.49 |
| A6208<sup>a,c</sup> | DP122A | MH560466 | *Cmn* | corn | NE, USA | 12.92 | 17.47 |
| A6209<sup>a,c</sup> | DP134 | MH560467 | *Cmn* | corn | TX, USA | 15.81 | 20.09 |
| A6210<sup>a,c</sup> | DP137 | MH560468 | *Cmn* | corn | TX, USA | 13.09 | 17.87 |
| A6211<sup>a,c</sup> | DP139B | MH560469 | *Cmn* | corn | TX, USA | 13.64 | 18.82 |
| A6212<sup>a,c</sup> | DP164 | MH560470 | *Cmn* | corn | NE, USA | 11.08 | 15.22 |
| A6213<sup>a,c</sup> | DP165c | MH560471 | *Cmn* | corn | IA, USA | 12.48 | 17.59 |
| A6094<sup>a,c</sup> | NCPPB 2579; LMG 3698 | MH560472 | *Cmn* | corn | NE, USA | 15.94 | 19.09 |
| A6095 <sup>a,c</sup> | 20037 | MH560473 | *Cmn* | corn | NE, USA | 10.06 | 14.91 |
| A6096 <sup>a,c</sup> | 200800460 | MH560474 | *Cmn* | corn | NE, USA | 12.05 | 16.84 |
| K-1 <sup>*,c</sup> |  | MH560475 | *Cmn* | corn | KA, USA | 11.71 | 15.65 |
| K-2 <sup>*,c</sup> |  | MH560476 | *Cmn* | corn | KA, USA | 10.01 | 14.71 |
| A2058 <sup>c</sup> | H-160 | MH560477 | *C. michiganensis* subsp. *michiganensis* (*Cmm*) | tomato | ID, USA | 17.5 | X |
| A4004 <sup>c</sup> | 47 | MH560481 | *Cmm* | tomato | OH, USA | 12.23 | X |
| A4763 <sup>c</sup> | N 7388A; K075 | MH560485 | *Cmm* | tomato | Morocco | 9.01 | X |
| A4758 <sup>c</sup> | N 212; K074 | MH560484 | *Cmm* | tomato | China | 8.86 | X |
| A4598 <sup>c</sup> | Cmm038 | MH560482 | *Cmm* | tomato | WA, USA | 10.71 | X |
| A4690 <sup>c</sup> | CMM 461 | MH560483 | *Cmm* | tomato | PORTUGAL | 10.01 | X |
| A2645 <sup>c</sup> | S47 | MH560480 | *Cmm* | tomato | CA, USA | 10.25 | X |
| A1746 <sup>c</sup> | A 518-1 | MH560479 | *Cmm* | tomato | HI, USA | 9.32 | X |
| A6268 <sup>c</sup> | 70049 | MH560492 | *Cmm* | tomato | Unknown | 16.88 | X |
| A2041 <sup>c</sup> | R8 | MH560493 | *C. michiganensis* subsp. *sepedonicus* | potato | Unknown | 19.08 | X |
| A6172 <sup>c</sup> | ATCC33113 | MH560494 | *C. michiganensis* subsp. *sepedonicus* | potato | Canada | 16.52 | X |
| A6101 <sup>c</sup> | ATCCBAA-2690; ZUM3936; CFBP 8217 | MH560495 | *C. michiganensis* subsp. *chiloensis* | tomato | Chile | 16.36 | X |
| A6112 <sup>c</sup> | 1646; P5005 | MH560497 | *C. michiganensis* subsp. *capsici* | pepper | China | 11.49 | X |
| A6113 <sup>c</sup> | 1647; P5006 | MH560498 | *C. michiganensis* subsp. *capsici* | pepper | China | 13.81 | X |
| A6134 <sup>c</sup> | C55 | MH560499 | *C. michiganensis* subsp. *californiensis* | tomato | CA, USA | 10.13 | X |
| A6135 <sup>c</sup> | LPPA982; LMG 27667 | MH560500 | *C. michiganensis* subsp. *phaseoli* | bean | Spain | 15.66 | X |
| A1149 <sup>c</sup> | QR-80 | MH560501 | *C. michiganensis* subsp. *insidiosus* | alfalfa | KS, USA | 16.5 | X |
| A6109 <sup>c</sup> | ZUM 4260; LMG7294; ATCC33566 | MH560502 | *C. michiganensis* subsp. *tessellarius* | wheat | USA | 14.8 | X |
| A1148 | QR-79; ATCC6887 | □ | *Curtobacterium flaccumfaciens* pv. *flaccumfaciens* | bean | Unknown | X | X |
| A1150 | QR-81a; ATCC12975 | □ | *Rhodococcus fascians* | unknown | USA | X | X |
| A1151 | QR-81b | □ | *R. fascians* | unknown | USA | X | X |
| A1147 | QR-78; ATCC9682 | □ | *Curtobacterium flaccumfaciens* pv. *poinsettiae* | poinsettia | USA | X | X |
| A6266 | 70002 | □ | *C. flaccumfaciens* pv. *poinsettiae* | poinsettia | Unknown | X | X |
| A6267 | 70008 | □ | *Corynebacterium flaccumfaciens* subsp. *oortii* | tulip | Unknown | X | X |
| A6272 | 70423 | □ | *C. flaccumfaciens* subsp. *oortii* | tulip | Unknown | X | X |
| A1152 | QR-82; ATCC13659 | □ □ □ | *Rathayibacter rathayi* | grass | United Kingdom | X | X |
| A6214 | DP101 | MH547375 | *Microbacterium* sp. | corn | IA, USA | X | X |
| A6218 <sup>e</sup> | DP122B | MH547376 | *C. flaccumfaciens* pv. *flaccumfaciens* | corn | NE, USA | X | X |
| A6216 <sup>e</sup> | DP114B | MH547381 | *Erwinia* sp. | corn | TX, USA | X | X |
| A6222<sup>e</sup> | DP138 | MH547382 | *Pantoea agglomerans* | corn | WI, USA | X | X |
| A6223<sup>e</sup> | DP140 | MH547379 | *Klebsiella* sp. | corn | IA, USA | X | X |
| A6225<sup>e</sup> | DP143 | MH547388 | *Pantoea ananatis* | corn | NM, USA | X | X |
| A6276<sup>e</sup> | DP150B | MH547383 | *P. agglomerans* | corn | MO, USA | X | X |
| A6220<sup>e</sup> | DP133 | MH547386 | *P. ananatis* | corn | IA, USA | X | X |
| A6227<sup>e</sup> | DP149 | MH547390 | *P. ananatis* | corn | TX, USA | X | X |
| A6224<sup>e</sup> | DP142 | MH547377 | *Pseudomonas* sp. | corn | IA, USA | X | X |
| A6230<sup>e</sup> | DP162A | MH547378 | *Pseudomonas* sp. | corn | SD, USA | X | X |
| A6234 | D36-1 | □ | *Xanthomonas axonopodis* pv. *dieffenbachiae* | anthurium | HI, USA | X | X |
| A5422 | CFBP2052 | □□ | *Dickeya zeae* | corn | USA | X | X |
| A5371 | CC26 | □ | *Pectovacterium carotovorum* | aglaonema | HI, USA | X | X |
| A3830 | CC46 | □□□ | *Pseudomonas syringae* pv. *syringae* | rice | South Africa | X | X |
| A6178 | CC36 | □ | *P. syringae* pv. *tomato* | tomato | USA | X | X |
| A3617 | CC93R | □ | *Xanthomonas vesicatoria* | tomato | South America | X | X |

<sup>a</sup> ATCC: American Type Culture Collection; CFBP: Collection Française de Bactéries associées aux plantes (French Collection of Plant associated bacteria); LMG: Collection of Laboratorium voor Microbiologie en Microbiële Genetica, Belgium; DP: strains isolated from corn leaf samples; <sup>b</sup> Average C<sub>T</sub> values calculated using Rotor-Gene Q Series Software (Qiagen Inc, Valencia, CA) using flap primers; <sup>c</sup> Strains used for corn leaf infection; <sup>d</sup> identity confirmation using *dnaA* gene region; identity confirmation using 16S *rRNA* gene region; <sup>\*\*</sup> extracted DNA from infected samples of corn was received from James P. Stack (Kansas State University-2017); <sup>e</sup> identity confirmation using 16S *rRNA* gene region.

### Sequencing and Identity Confirmation

A new primer set P16s-F1 (5’-AGACTCCTACGGGAGGCAGCA-3’) and P16s-R1 (5’- TTGACGTCATCCCCACCTTCC-3’) targeting the 16S ribosomal RNA region (16S *rRNA*) of plant bacteria was designed and used to identify eleven isolates from corn leaves that showed characteristic symptoms of Goss’s disease; the strains were provided by DuPont Pioneer Plant Disease Diagnostic Laboratory during the 2015 growing season (“DP” number; S1 Table). Also, a new set of primers CM-dnaA-F1 (5’-ACGAAGTACGGCTTCGACAC-3’) and CM-dnaA-R1 (5’-GCGGTGTGGTTGATGATGTC-3’) was designed to amplify a partial sequence of the chromosomal replication initiator gene, *dnaA*, of *C. michiganensis*. PCR assays for 16S rRNA and *dnaA* were performed with the following conditions: initial denaturation at 94°C for 5 min, 35 cycles of denaturation at 94°C for 20 sec, annealing at 58°C for 30 sec, extension at 72°C for 1 min, and a final extension at 72°C for 3 min. Five μl of PCR product was cleaned enzymatically using 2 μL of ExoSAP-IT™ following the manufacturer’s protocol (Affymetrix Inc, Santa Clara, CA). Sanger sequencing of both sense and anti-sense strands was performed at GENEWIZ facility (Genewiz, La Jolla, CA). Obtained sense and anti-sense sequences were aligned and manually edited for accuracy using Geneious version 10.1.3. Consensus sequences were compared against the NCBI GenBank nucleotide and genome databases using BLASTn tool [20].

### Phylogenetic Analysis

A total of forty-three partial sequences belonging to the Chromosomal initiation factor gene, *dnaA*, were used to perform the phylogenetic analysis. Generated sequences from thirty-two strains of *C. michiganensis* were submitted to NCBI GenBank nucleotide database (S1 Table). DNA sequences from ten Gram-positive plant bacteria were retrieved from GenBank. Accession numbers pertain to nine *C. michiganensis* species (JX259295.1, HE614873.1 and KF663898.1 for *C. m.* subsp. *nebraskensis*; CP011043.1 for *C. m.* subsp. *insidiosus*; CP0125523.1 for *C. m.* subsp. *capsici*; CP012573.1 and KJ723744.1 for *C. m.* subsp. *michiganensis*; and AM849034.1 for *C. m.* subsp. *sepedonicus*), and two sequences from relatives to the closest taxa (CP014761.1, *Leifsonia xyli;* CP015515.1, *R. tritici*), which also served as the outgroup.

All the sequences were aligned using muscle algorithm [21] plugged in MEGA 7.0.25 [22]. For single gene *dnaA*, the best evolution model was determined, and maximum likelihood (ML) method with 1,000 bootstrap values to evaluate node support was used to infer the phylogenetic tree. A color-coded matrix showing pairwise identity was generated using Sequence Demarcation Tool version 1.2 [23].

### Infected Plant Materials and DNA Isolation

Seeds of hybrid sweet corn variety Hawaiian supersweet #10 were provided by UH seed lab, HI (Agricultural Diagnosis Service Center); seeds were sown in Sunshine#4 soil mix in 20 cm pots (1 seed/pot) – experiment was conducted in 2017 at Pope Greenhouse, College of Tropical Agriculture and Human Resources, University of Hawaii at Manoa. Fourteen strains of *C. m.* subsp. *nebraskensis* (S1 Table) were grown on YSC medium (yeast extract 10 g/L, sucrose 20 g/L, calcium carbonate 20 g/L and agar 17 g/L) and incubated at 26°C (±2°C) for 3 days; bacterial cells were harvested by flooding each plate with 5 ml of PBS buffer (pH 7.0). Corn plants (V-3 developmental stage) were inoculated using the dropped-wounded protocol described by Ahmad and collaborators [24] with a minor modification - the third leaf of each plant was wounded half way between the sheath and the tip by making a cut across 3 veins on one side of the midrib; immediately after wounding, 10 μl of inoculum was dropped on to the wound using a plastic dropper. Negative control plants were inoculated with 10 μl PBS (mock inoculated). Inoculated plants were covered with a plastic bag for 24 hours to increase the humidity to facilitate infection. Leaf samples from inoculated plants were collected 30 days post inoculation (dpi); DNA was isolated from each sample using the Wizard^®^ Genomic DNA Purification Kit according to the manufacturer’s instructions.

### Comparative Genomic Analysis for Target Selection and Primers/Probes Design

Potential gene targets for broad range and specific detection of *C. michiganensis* subspecies and *C. m.* subsp. *nebraskensis*, in particular, were identified by comparative genomic analysis using Mauve [25] and Geneious. To calculate the core genome in the *C. michiganensis* subspecies and the variable genomes for each subspecies, the annotated genomes of gram-positive bacteria, *C. m.* subsp. *nebraskensis* NCPPC 2581 (NC_020891), *C. m.* subsp. *michiganensis* NCPPB 382 (NC_009480), *C. m.* subsp. *sepedonicus* ATCC 33113 (NC_010407), *C. m.* subsp. *insidiosus* strain R1-1 (NZ_CP011043), and *C. m.* subsp. *capsici* strain PF008 (NZ_CP012573) were used. Moreover, *Rathayibacter tritici* (NZ_CP015515), *R. toxicus* (NZ_CP013292), *Rhodococcus fascians* (CP015235), *Corynebacterium efficiens* (CP004369), and *Curtobacterium* sp. (CP009755) genomes were included to confirm primer specificity developed in this study. Manually filtered orthologous genes present in the core genome of the five subspecies were identified as potential targets for detection of all known nine *C. michiganensis* subspecies. Putative and characterized genes in the accessory genome of *C. m.* subsp. *nebraskensis*, no homology with genes from other *C. michiganensis* subspecies, were selected as possible targets to detect *C. m.* subsp. *nebraskensis*. Homology and specificity of the *C. michiganensis*-specific (ABC transporter ATP-binding protein CDS/ABC-transporter permease CDS) and *C. m.* subsp. *nebraskensis*-specific (Major facilitator superfamily (MFS) transporter CDS) candidate genes were validated *in-silico* using NCBI GenBank BLASTn analysis tool. The probe CM-P and Cmn11-P were labeled with Cy5 and FAM reporter dyes, respectively. All primers and probes were designed using Primer3; thermodynamic parameters, internal structures, and self-dimer formation were evaluated *in silico* Primer3 and mFold web server (Table 2). Primers and probes were synthesized by Integrated DNA technologies Inc. (Coralville, IA).

**Fig 1.**
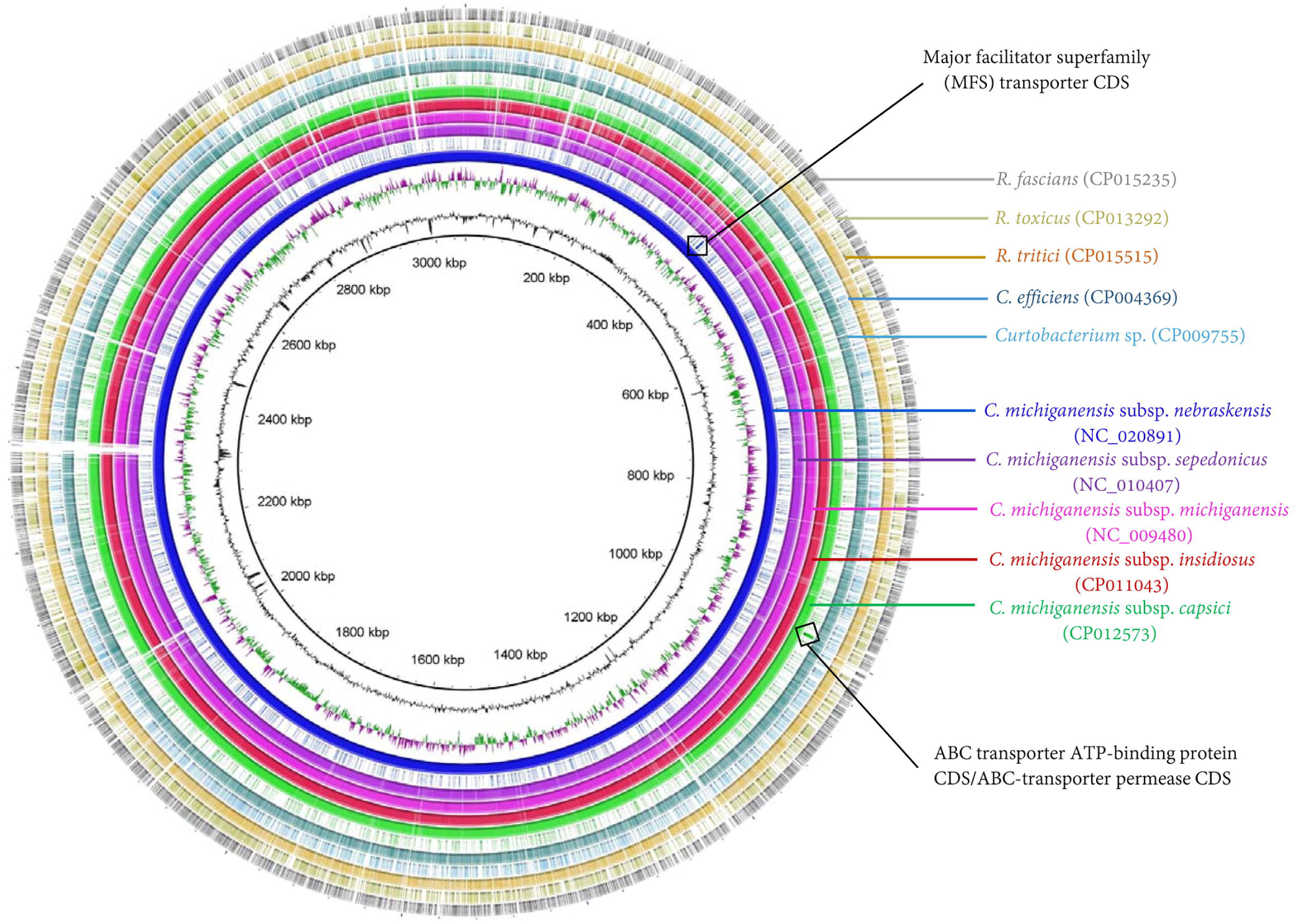
Genome alignment, comparison and locations of target genes unique to *C. michiganensis* and *C. michiganensis* subsp. *nebraskensis.* A ring image was generated to locate ABC transporter ATP-binding protein/ABC-transporter permease and Major facilitator superfamily (MFS) transporter gene regions for *C. michiganensis* and *C. michiganensis* subsp. *nebraskensis*, respectively. Strains used to generate a BRIG image are mentioned in the figure with their NCBI GenBank accession numbers in parenthesis; all the genomes were retrieved from NCBI GenBank genome database. The mapped genome ring image from the inside out shows: genome coordinates (kbp), GC content (black), GC skew (purple/green). The remaining rings show BLASTN comparison of 10 complete genomes and location of target genes following as labelled. *Clavibacter michiganensis* subsp. *nebraskensis* (NC_020891) was used as reference genome.

**Table 2.**
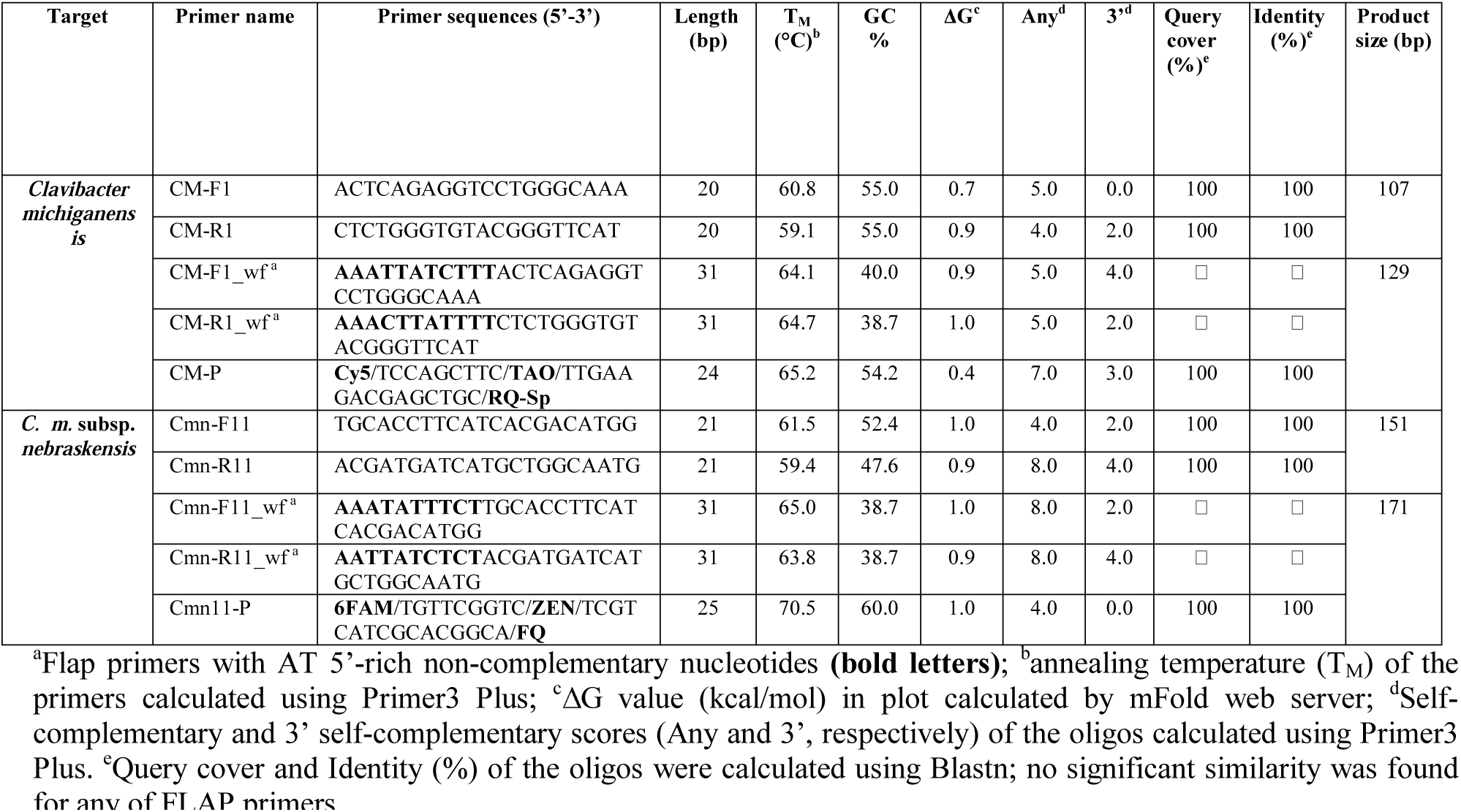
Thermodynamic parameters, secondary structures and product size of the primers and probes designed to specifically detect *Clavibacter michiganensis* and *C. michiganensis* subsp. *nebraskensis*.

### Addition of 5’ AT-rich Flap to Enhance the Reaction Efficiency

To evaluate the potential impact of 5’ AT-rich flap sequences, additional primer sets were synthesized after incorporating a customized 5’ AT-rich flap as described by Arif and Ochoa-Corona [17] (Table 2; Fig 2). The customized flaps were designed to finely tune the tm, primer length, and GC content to optimize the reaction efficiency. Eleven and ten nucleotide long flaps were customized and added to the primer sets for *C. michiganensis* and *C. m.* subsp. *nebraskensis*, respectively. No 5’ AT-rich flap sequences were added to the probes.

**Fig 2.**
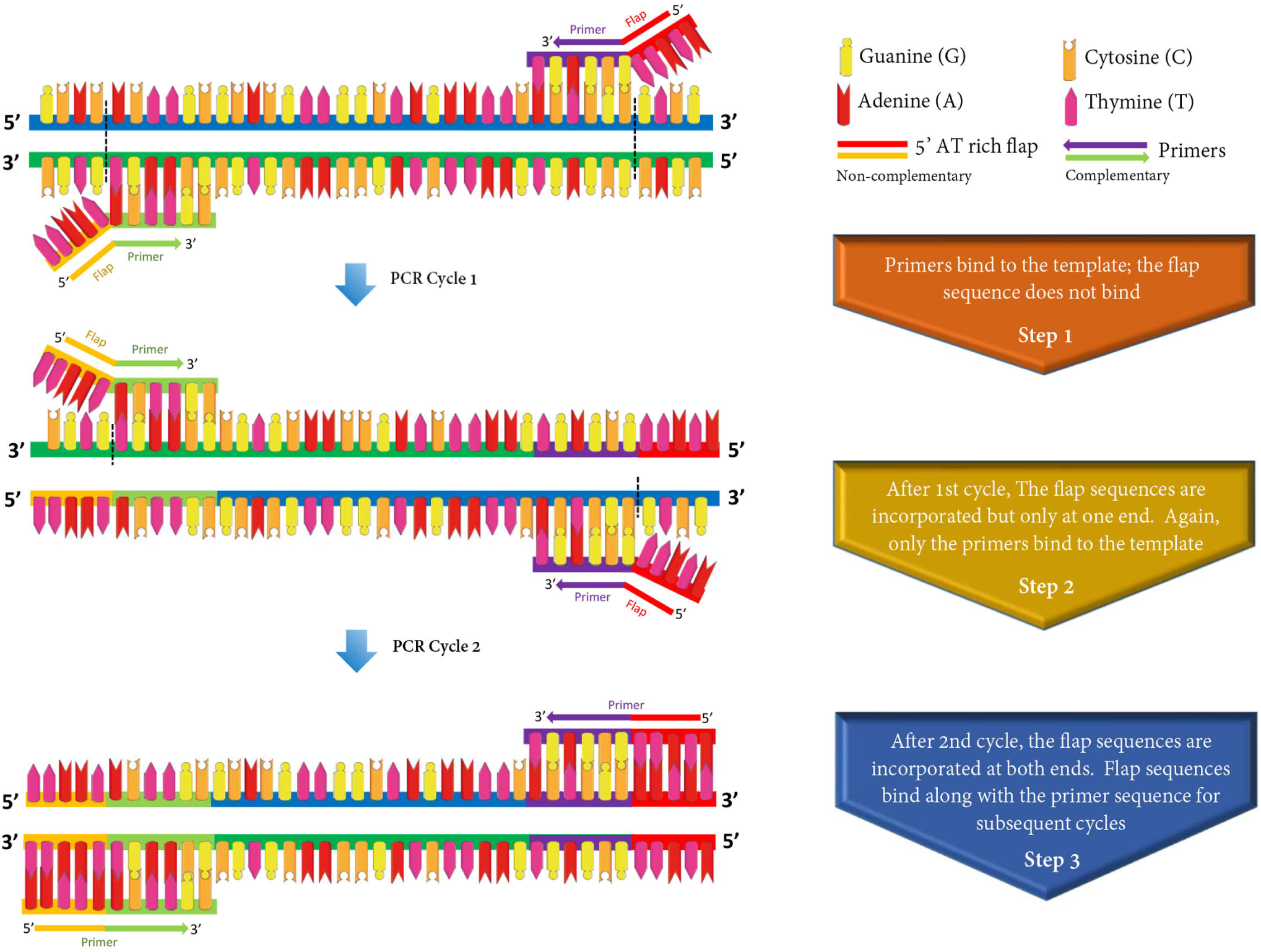
Principle and application of 5’-AT rich flap sequence to optimize polymerase chain reaction (PCR) thermodynamics and efficiency. The thermodynamics of both forward and reverse primers was improved by adding customized 10-11 nucleotides long AT rich sequences at 5’ position of each primer. The flap sequences are incorporated after the 2^nd^ cycle of the PCR – increases the specificity and speed with optimized reaction efficiency.

### Single and Multiplex End-point PCR

Individual single reactions to assess the four sets of primers for *C. michiganensis* subspecies and *C. m.* subspecies *nebraskensis* were carried out in 20 μl reactions containing 10 μl of 2X GoTaq® Green Master Mix (Promega), 1 μl of each forward and reverse primer (5 μM), 0.6 μl of MgCl2 (10 mM), 1 μl of DNA template, and 6.4 μl of nuclease-free water (VWR, Radnor, PA). The cycling parameters were: initial denaturation at 95°C for 5 min, followed by 35 cycles of denaturation at 95°C for 30 secs, annealing at 60°C for 30 secs, extension at 72°C for 30 secs, and a final extension at 72°C for 3 min. Multiplex assays for simultaneous detection of all nine subspecies within *C. michiganensis* and specific detection of *C. m. subspecies nebraskensis* were performed in a volume of 20 μl containing 10 μl of 2X GoTaq® Green Master Mix, 1 μl of each forward and reverse primer (5 μM), 0.6 μl of MgCl2 (10 mM), 1 μl of DNA template, and 4.4 μl of nuclease-free water. Thermal cycling parameters were the same as for the individual single reactions, but the annealing time was increased to 60 secs. PCR assays were conducted in a T100(tm) BioRad thermal cycler (Bio-Rad, Hercules, CA). Non-template controls were included in each PCR amplification - 1 μl of nuclease-free water was used instead of genomic DNA. A volume of 13 μl of amplified PCR product was used for electrophoresis in a 3% agarose gel; 1X Tris-acetate-EDTA (TAE) buffer at 80 V for 90 min was used for electrophoresis. Amplicon sizes were estimated using 100 bp DNA ladder (New England Biolabs, Ipswich, MA). Genomic DNA of *C. m.* subsp. *nebraskensis* strain A6206 was used to perform the PCR sensitivity assays with primer sets CM-F/R and Cmn11-F/R.

### Single and Multiplex TaqMan Real-time qPCR Assays

Multiplex TaqMan real-time qPCR assays were conducted in a volume of 25 μl, containing 12.5 μl of 2X Rotor-Gene® Multiplex PCR Master Mix (Qiagen, Valencia, CA), 2 μl of primer mix (6.25 mM), 0.5 μl of CM-P (5 mM), 0.5 μl of Cmn11-P (5 mM), 8.5 μl of nuclease-free water and 1 μl of DNA. Single qPCR assays were performed in a reaction volume of 25 μl containing 12.5 μ of 2X Rotor-Gene® Multiplex PCR Master Mix, 2 μl of CM or Cmn primer mix reaction (6.25 mM), 0.5 μl of probe (5 mM), 9 μl of nuclease-free water and 1 μl of DNA. CM and Cmn primer sets were added to a final concentration of 0.5 µM for each primer in single and multiplex TaqMan qPCR assays. Likewise, final concentrations of 0.2 µM for each probe and 3 mM of MgCl_2_ were used in single and multiplex assays. Each TaqMan qPCR reaction was conducted in three replicates. Rotor-Gene Q (Qiagen) was used to perform the TaqMan qPCR assays. The cycling conditions consisted of a PCR initial activation step at 95°C for 5 min, following by rapid 40 cycles of denaturation at 95°C for 15 sec and annealing/extension at 60°C for 15 sec. Cycle threshold (C_T_) values were calculated and analyzed with Rotor-Gene® Q series software (Qiagen).

### End-point, TaqMan qPCR Sensitivity and Spiked Assays

To determine sensitivity and PCR yields, multiplex end-point PCR and TaqMan real-time qPCR assays were performed employing two sets of primers with and without 5’ AT-rich sequences (Table 2). Sensitivities were determined using 10-fold serially diluted genomic DNA; the DNA concentrations ranged from 10 ng to 10 fg per reaction. Pure bacterial DNA from *C. m.* subsp. *nebraskensis* (A6206) was used for all sensitivity assays; NanoDrop v.2000 spectrophotometer (Thermo Fisher Scientific, Inc., Worcester, MA) was used to measure DNA concentrations. All TaqMan qPCR reactions were performed in three replicates. The spiked assays were performed by adding 1 µl of healthy corn host DNA into TaqMan qPCR sensitivity reactions prepared using serially diluted (10 ng to 10 fg) genomic DNA of *C. m.* subsp. *nebraskensis* (A6206).

## Results

### Sequencing and Identity Confirmation

Two sets of primers targeting two housekeeping genes, 16S *rRNA* for all plant-associated bacteria and *dnaA*-specific for *C. michiganensis* subspecies, resulted in species and subspecies identification, respectively. Eleven bacterial strains isolated from corn were identified using the novel P16S-F/R primer set resulting in an 830-870 bp amplicon. Two gram-positive bacteria were identified as *Curtobacterium* sp. (A6218) and *Microbacterium* sp. (A6214); also, nine Gram-negative bacteria were identified as *Erwinia* sp., *Klebsiella* sp., *Pantoea* sp. and *Pseudomonas* sp. by 16S *rRNA* gene sequence (S1 Table). Strains A6214 and A6218 displayed similar colony morphologies as *C. michiganensis* when cultured on TCZ-S medium (data not shown). Identity of fourteen *C. michiganensis* subsp. *nebraskensis* strains isolated from corn and two DNA samples from corn plants showing Goss’s symptoms (J. Stack, Kansas State University, 2017) were confirmed in the first test as subspecies within *C. michiganensis* using CM-dnaA-F1/R1 primer set. Moreover, identity of eighteen strains within the remaining eight subspecies of *C. michiganensis* isolated from different hosts and geographical locations, was confirmed using the same CM-dnaA-F1/R1 primer set (S1 Table).

The amplicon generated using either P16S or CM-dnaA primer sets exhibited 100% query coverage and 100% identity against the sequence corresponding to species/subspecies sequence present in NCBI the nucleotide and genome databases, suggesting that the novel primer sets are able to accurately identify plant-associated bacteria as well as the nine subspecies within *C. michiganensis*.

### Sequence Alignment and Phylogenetic Analyses

The General Time Reversible with gamma distribution model (GTR + G) was used to construct the maximum likelihood tree. The different subspecies within *C. michiganensis* were grouped in three main clusters. *C. m.* subsp. *tessellarius* was grouped discretely in one clade. In contrast, the remaining eight subspecies were placed into two clades. In the first clade, *C. m*. subsp. *nebraskensis, C. m*. subsp. *insidiosus, C. m*. subsp. *capsici, C. m*. subsp. *phaseoli* and *C. m*. subsp. *chiloensis* were grouped. Interestingly, isolates of *C. m.* subsp. *nebraskensis* were placed together with *C. m*. subsp. *insidiosus* displaying 97% of similarity, while two subspecies, *C. m*. subsp. *phaseoli* and *C. m.* subsp. *chiloensis*, were placed together in a single branch and simultaneously grouped with *C. m*. subsp. *capsici* sharing 76% of similarity. The second clade grouped *C. m*. subsp. *michiganensis, C. m.* subsp. *californiensis* and *C. m*. subsp. *sepedonicus*. *C. m*. subsp. *michiganensis* strains were highly related to *C. m.* subsp. *californiensis* with 99% of similarity (Fig 3). Additionally, the axis on the color-coded matrix confirmed the diversity within *C. michiganensis* subspecies ranging from 89 to 98% (Fig 3).

**Fig 3.**
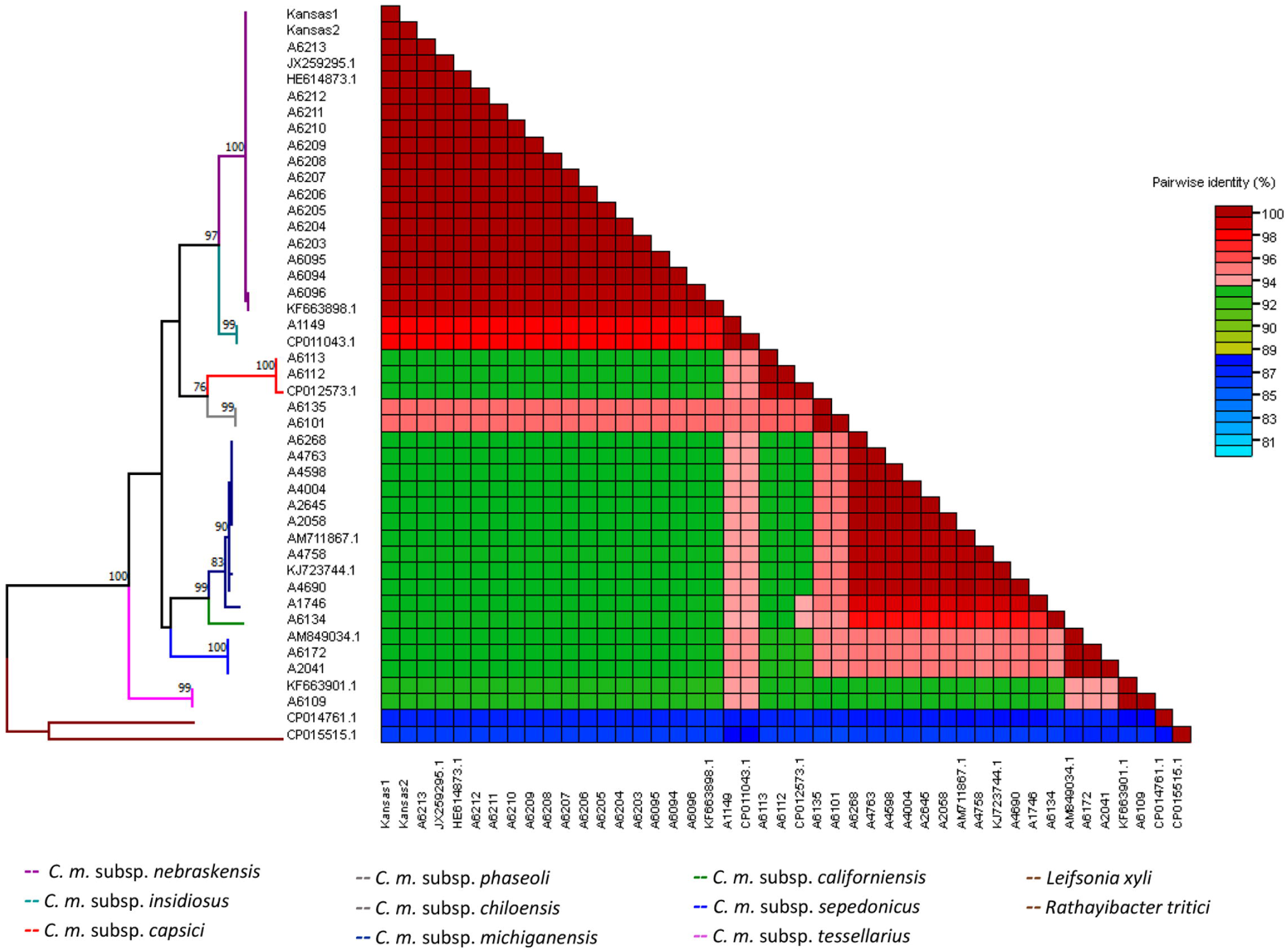
*dnaA* color-coded demarcation matrix of *Clavibacter michiganensis* subspecies. maximum likelihood (ML) phylogeny of 43 partial sequences within *C. michiganensis* subpecies were used for constructing the pairwise similarity scores, displaying a color-coded matrix. *dnaA* phylogeny within *C. michiganensis* shows evolutionary distances gathering nine evident clades, each one representing one subspecies. Bootstrap values were computed with 1,000 replicates using MEGA 7.0.25 and *dnaA* sequences of *Leifsonia xyli* and *Rathayibacter triciti* served as outgroup, sharing less than 89% similarity.

### Target Selection and Primer Design

Highly conserved regions within the major facilitator superfamily (MFS) transporter gene for *C. michiganensis* subsp. *nebraskensis*, ABC transporter ATP-binding protein CDS/ABC-transporter permease genes for *C. michiganensis* subspecies were used to design two sets of primers for both conventional end-point PCR and TaqMan real-time qPCR (Table 2; S2 Fig). The primers and probes evaluated *in silico* showed high specificity for *C. m.* subsp. *nebraskensis* (Cmn) and *C. michiganensis* subspecies in general (Table 2). No nucleic acid similarity to orthologous genes or conserved domains were found for the major facilitator superfamily (MFS) supporting the unique presence of MFS gene only in *C. michiganensis* subsp. *nebraskensis*. On the other hand, highly conserved domains of sugar ABC transporter permease and ABC transporter ATP-binding genes within *C. michiganensis* subspecies were selected to design specific primers and probe, resulting in a broad but specific detection of all nine subspecies within *C. michiganensis*. Primer and probe sets were highly specific as revealed by NCBI BLASTn analysis.

### *In vitro* Specificity of the Developed Assays

Each primer and probe set (Table 2) was designed to be used in both endpoint PCR (endpoint PCR products between 100 to 200 bp) and TaqMan real-time qPCR (both multiplex and single reaction). Broad range detection capabilities and specificity of primers and probes were tested against inclusivity and exclusivity panels containing twenty-nine strains of *C. michiganensis* subspecies and twenty-five gram-positive and gram-negative bacteria from either symptomatic corn leaf or closely related plant associated bacteria (S1 Table). Fourteen strains and two naturally infected samples were unmistakably detected as *C. michiganensis* subsp. *nebraskensis* using primer Cmn11-F/R and probe Cmn11-P; no cross-reactivity was observed with any other non-target species/subspecies (Table 1). Likewise, the primer (CM-F/R) and probe (CM-P) set exhibited an accurate and specific detection of all twenty-nine strains corresponding to the nine subspecies of *C. michiganensis*; no cross-reactivity with the members of exclusivity panel was detected. As expected, 151-and 107-bp PCR amplicons were amplified in multiplex endpoint PCR for *C. michiganensis* subsp. *nebraskensis* and *Clavibacter michiganensis* species, respectively (S3 Fig).

### Effect of AT 5’-rich Flaps on the Sensitivity and Reaction Efficiency

All primers and probes met the desired 100% query coverage and 100% identity after an alignment using BLASTn against the NCBI nucleotide and genome databases (Table 2). Thermodynamics, internal structures, GC% content and ΔG values for each primer and probe were calculated using mFold and Primer3 Plus; all the parameters are listed in Table 2. The tendency of primers to form self-structures increased when additional nucleotides were added to the primers; nevertheless, no interference in primer annealing with the target sequence was observed when 5’ AT-rich sequences were added to the primers. Furthermore, assay sensitivity for primer sets with flaps was increased ten-fold suggesting positive impact on reaction thermodynamics and stability during annealing and extension steps (Fig 2).

The detection limits of TaqMan real-time qPCR single reactions for both primer and probe sets were 100 fg using purified bacterial genomic DNA (Fig 4A and B). However, low reaction efficiencies of 82% and 91% were observed for *C. michiganensis* species and *C. m.* subsp. *nebraskensis* targets, respectively, suggesting inhibition or sub-optimal conditions for amplification (Fig 4A-B Table 3). To optimize the amplification conditions, customized 5’AT-rich sequences were added to each primer; the addition of AT-rich sequences at the 5’ position remarkably enhanced the reaction efficiency to 99-104% (Fig 4C-H). The most pronounced positive effect was observed with the *C. michiganensis* specific primer and probe set; the sensitivity was enhanced 10-fold and detected the target sequence in 10 fg of genomic DNA (Fig 4A and B; Table 3). No adverse effect was observed when 1 µl host plant DNA was added in each sensitivity reaction (spiked test, Fig 4E). Similarly, no effect was observed with the *C. m.* subsp. *nebraskensis*-specific primers and probe designed using the MFS gene region. The AT-rich sequence effect was also evaluated with endpoint PCR, the *C. michiganensis* assay sensitivity was enhanced from 100 fg to 10 fg (S3 Fig). Additionally, analysis of the amplification curves indicated that the use of 5’AT-rich sequences did not affect the sigmoidal shape with the concentration of target in sensitivity assays. In contrast, use of non-tailed primers negatively impacted the amplifications as well as the efficiencies (Fig 4).

**Table 3.**
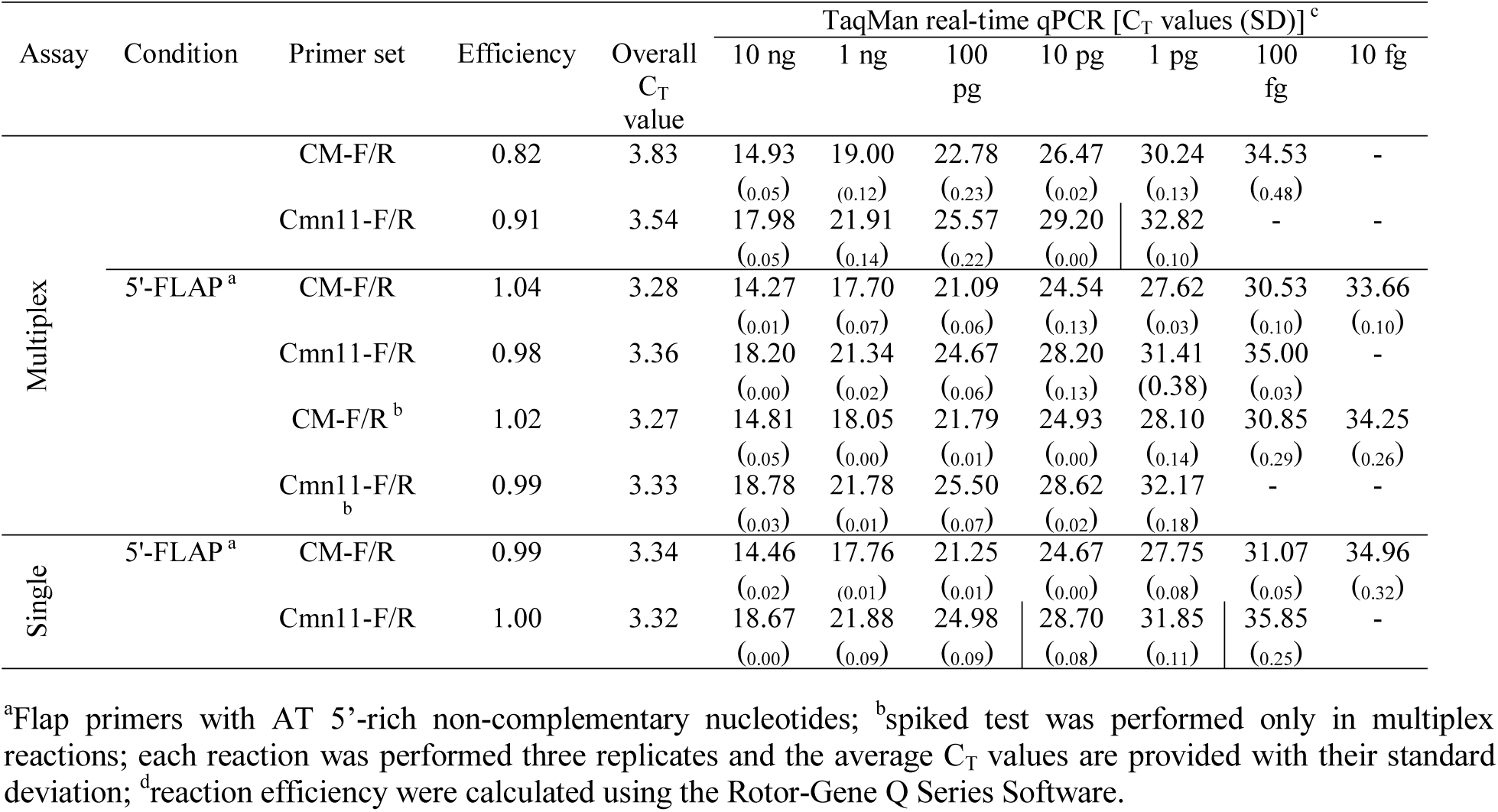
Comparative analyses of 5’ AT-rich tailed and non-tailed primers specifically designed to detect *Clavibacter michiganensis* and *C. michiganensis* subsp. *nebraskensis.*

**Fig 4.**
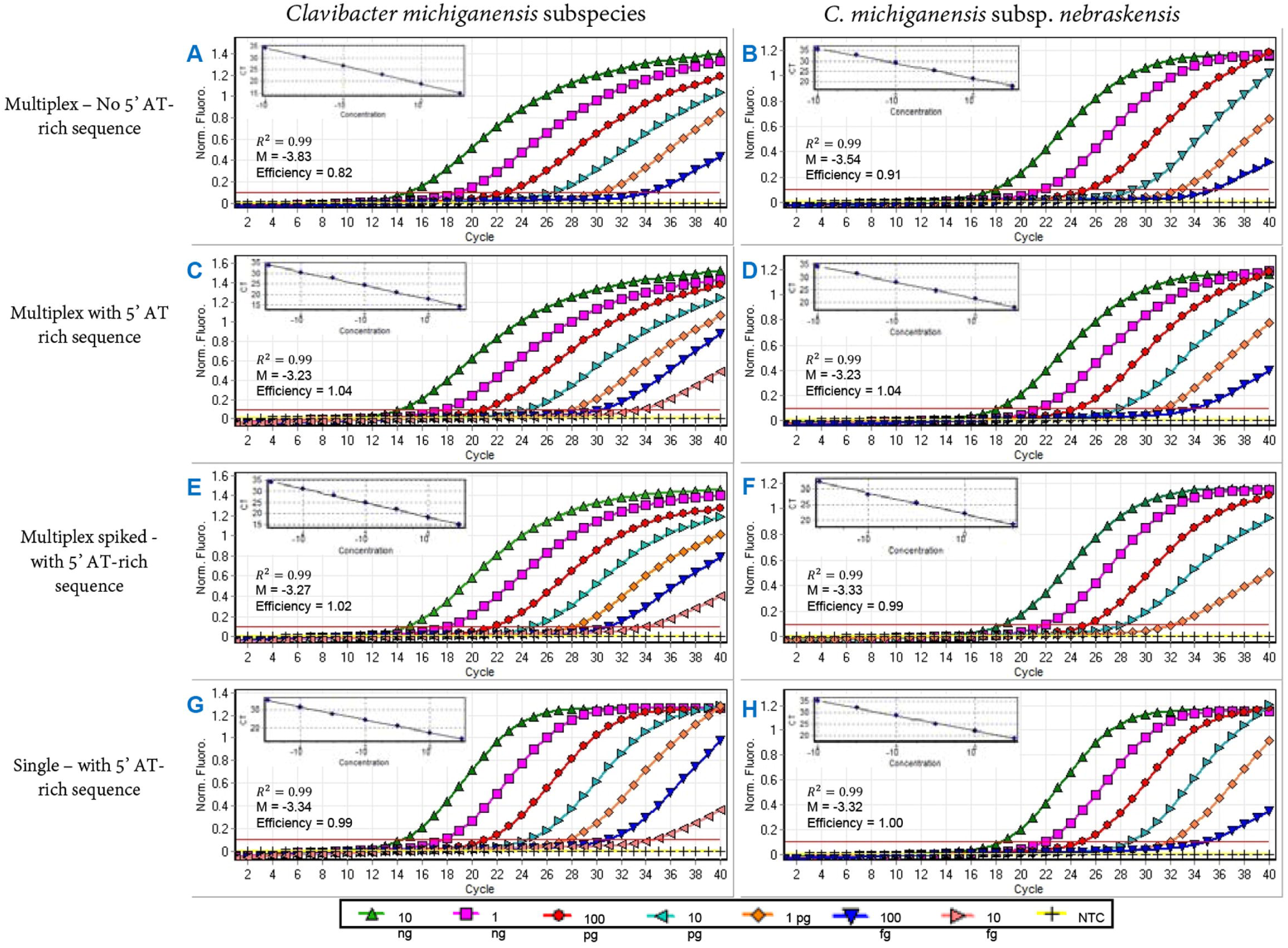
Effect of 5’ AT-rich flap on TaqMan real-time qPCR sensitivity assays performed using ten-fold serially diluted genomic DNA. **(A)** and (**B)**. TaqMan real-time qPCR assays with individual primer set harboring no complementary 5’ AT-rich sequences. (**A)** represents the detection *C. michiganens*is; and **(B)** represents the subspecific detection of *C. m.* subsp. *nebraskensis*; (**C, D, E, F** and **G**) are multiplex and single reactions after adding the non-complementary 5’ AT-rich sequences. **(C)** and **(D)** multiplex reactions for simultaneous detection of all subspecies of *C. michiganens*is and of *C. m.* subsp. *nebraskensis*, respectively. **(E)** and **(F)** multiplex reactions conducted for simultaneous detection of *C. michiganens*is subspecies and of *C. m.* subsp. *nebraskensis*, respectively, after adding 1 μl of healthy plant corn DNA to each 10-fold serially diluted sensitivity reactions in order to simulate natural infection. **(G)** and **(H)** TaqMan real-time qPCR assays performed with individual primer and probe sets to detect *C. michiganens*is subspecies and *C. m.* subsp. *nebraskensis*, respectively. All assays were conducted from 10 ng to 10 fg serially diluted genomic DNA of *C. m.* subsp. *nebraskensis* (strain A6206) using Rotorgene.

## Discussion

We describe the enhancing effect of 5’ AT-rich sequences on reaction efficiency with fast and rapid TaqMan qPCR cycling conditions. The primers and probes were designed using unique and conserved gene regions to detect *C. michiganensis* species in general as well as specifically, *C. m.* subsp. *nebraskensis*. The developed assays are highly sensitive and specific.

Population genetic and comparative genomic analyses are imperative components for the development of robust and specific diagnostic tools (5,26,27,28). Multiple genome alignments using Mauve software resulted in highly conserved and unique gene regions specific for *C. michiganensis* in general (major facilitator superfamily gene) and specifically for *C. m.* subsp. *nebraskensis* (core sugar ABC transporter permease and ABC transporter ATP-binding genes). Besides the selection of unique and conserved targets, primer and probe design plays a very important role in developing a robust and specific multiplex TaqMan qPCR assay [17,29]. Assay specificity can also be enhanced using dual-quenched probes [16,30]; that is, an internal quencher, TAO or ZEN, was added to each probe to reduce the distance from reporter to quencher, leading to low background signal and greater precision (Table 2).

Arif and Ochoa-Corona (2013) [17] found that the addition of non-complementary AT-rich sequences at the 5’ position of the forward and reverse primers enhanced the detection sensitivity and overall PCR amplicon yields; they reported that the addition of customized sequences on suboptimal (thermodynamically incompetent) primers increased the PCR yield by about 20% and sensitivity by 10-100 folds. In the current study, we observed that the addition of 5’ AT-rich sequences not only increased the sensitivity but also optimized the reaction efficiency from 82% to 99% and 91% to 100% for *C. michiganensis* in general and *C. michiganensis* subsp. *nebraskensis*, respectively (Fig 4). One hypothesis is that addition of 5’ AT-rich sequences to the primers enhanced the reaction thermodynamics allowing for faster primer annealing to the target template resulting in optimal reaction efficiency; however, the exact mechanism is unknown. The outcome of flap addition also resulted in a 10-fold increase in sensitivity for detection of *C. michiganensis* (Fig 4; Table 3). Although the overall mechanism involved in improved sensitivity and efficiency due to the use of flap primers in TaqMan qPCR assays is still not completely clear, a study conducted by Vandenbussche and group (2016) [31] suggests that tailed primers suppress the formation of PCR artifacts, determined by primers and dNTP type. Also, tailed primer reactions respond differently to the use of hot-start dNTPs. In this study, the use of the TaqMan real-time qPCR master mix containing hot-start dNTP led to specific detection and amplification without the report of either PCR artifacts or shifting in the sigmoidal shape of the amplification curves.

The developed TaqMan qPCR assays have proven to be truly sensitive and efficient for simultaneous and single detection of *C. michiganensis* and *C. m.* subsp. *nebraskensis*. The assays were fully validated with well-described strains in inclusivity and exclusivity panels. The universal primer set P16-F/R designed to amplify the 16S rRNA partial gene region of plant-associated bacteria provided genus and species identification for the eleven bacterial pathogens from corn, including gram-positive and gram-negative bacteria (S1 Table). Furthermore, primer set CM-dnaA-F/R, designed to amplify the *dnaA* partial-gene sequence of *C. michiganensis*, was used to confirm the identity of all nine subspecies (S1 Table). The housekeeping *dnaA* gene has been proven as a potential genetic marker for plant bacterial phylogenetic studies and identity confirmation [5].

Two new non-pathogenic subspecies, *C. m.* subsp. *californiensis* and *C. m.* subsp. *chiloensis*, associated with tomato as a common host [2] but only *C. m*. subsp. *californiensis* clustered with *C. m*. subsp. *michiganensis*. In contrast, *C. m.* subsp. *chiloensis* clustered with *C. m*. subsp. *phaseoli* (infects bean) and formed the group with *C. m.* subsp. *capsici* (infects pepper). Additionally, *C. m.* subsp. *nebraskensis* (infects corn) was grouped in the same clade as *C. m.* subsp. *insidiosus* (infects alfalfa). Interestingly, *C. m*. subsp. *tessellarius* formed a separate cluster closest to *C. m.* subsp. *nebraskensis* (Fig 3). The same phenomenon was observed when the FASTA pairwise alignment was plotted in the color-coded matrix (Fig 3). The current phylogenetic analysis using *dnaA* gene is congruent with previously reported studies using *gyr*B, *rec*A, *rpo*B, *gapA, icdA, mdh, mtl*D, *pgi* and *pro*A genes supporting the new taxonomy and classification of *Clavibacter* species [1,5].

Accurate detection and identification are critical in diagnosis and microbial forensics for disease control and management. The use of a robust multigene format maximizes broad-range detection capabilities, reliability and specificity for *C. michiganensis* species detection. Fifty-nine strains representing different diseases and hosts were included in the inclusivity and exclusivity panels – no false positives or false negatives were detected. Of 59 samples, 29 were identified as members of various *C. michiganensis* subspecies, 16 were identified as *C. m.* subsp. *nebraskensis* and 25 samples (other than *Cm* and *Cmn*) showed no reactivity when specific TaqMan qPCR assays were performed (Table 2). The positive qPCR results with genomic DNA of twenty-nine diverse strains in the inclusivity/exclusivity panels collected from different geographical locations across USA and the world confirmed specificity and broad-range identification of *C. michiganensis* subspecies and *C. m.* subsp. *nebraskensis* using the novel primer and probe sets developed in the present study (Table 1).

Multiplex TaqMan qPCR assays for detection of *C. michiganensis* species and *C. m.* subsp. *nebraskensis* are robust, accurate, sensitive and reliable. The developed assays can be used to accelerate phytosanitary diagnostics and target pathogen detection. Sensitive detection can help in early detection of latently-infected plant materials and support further studies in pathogen dissemination during interstate or international commerce. The addition of 5’ AT-rich sequences lead to the development of thermodynamically competent primers for sensitive detection. This concept also will enable development of improved protocols for different pathogens that can work under the same PCR conditions. Development of mix and match diagnostic protocols – where the primers can be multiplexed based on individual needs will be possible since each primer’s thermodynamics, GC content, annealing temperature and primer length can be customized to fit existing PCR conditions.

## Acknowledgments

This work was supported by the USDA National Institute of Food and Agriculture, Hatch project 9038H, managed by the College of Tropical Agriculture and Human Resources, and National Science Foundation [NSF; Award Number – 1561663]. The mention of trade names or commercial products in this publication does not imply recommendation or endorsement by the University of Hawaii. We thank to Dr. Jarred Yasuhara-Bell (Kansas State University) for isolating DNA from two naturally infected corn samples. The funders had no role in study design, data collection and analysis, decision to publish, or preparation of the manuscript.

## Author Contributions

**Conceptualization:** Mohammad Arif.

**Data curation:** Adriana Larrea-Sarmiento, Anne M. Alvarez, Mohammad Arif.

**Formal analysis:** Adriana Larrea-Sarmiento, Mohammad Arif.

**Funding acquisition:** Mohammad Arif, Anne M. Alvarez.

**Investigation:** Adriana Larrea-Sarmiento, Mohammad Arif.

**Methodology:** Adriana Larrea-Sarmiento, Mohammad Arif.

**Project Administration:** Mohammad Arif.

**Resources**: Anne M. Alvarez, Mohammad Arif.

**Supervision:** Mohammad Arif.

**Validation:** Adriana Larrea-Sarmiento, Mohammad Arif

**Visualization:** Adriana Larrea-Sarmiento, Mohammad Arif.

**Writing – original draft:** Adriana Larrea-Sarmiento

**Writing – review & editing:** Adriana Larrea-Sarmiento, Anne M. Alvarez, James P. Stack, Mohammad Arif.

## Supporting information captions

**S1 Fig.** *dnaA* phylogeny within the nine *C. michiganensis* subspecies inferred using the maximum likelihood (ML) method in MEGA 7.0.25. The evolutionary distances were computed using a General Time Reversible model with gamma distribution (GTR + G). Forty-three partial sequences based on *dnaA* gene of *Clavibacter* strains were rooted with 2 closest taxa *Leifsonia xyli* and *Rathayibacter triciti*, served as an outgroup. A 1,000 replicates were performed to calculate the bootstrap value which is shown over the branches; only bootstrap values greater than 70% are presented.

**S2 Fig.** Target selection. A. Primers and probe (Cmn11-F/R/P) targeting the MFS transporter gene specific for *C. m.* subsp. *nebraskensis.* B. Primers and probe (CM-F/R/P) targeting the sugar ABC transporter permease and ABC transporter ATP-binding genes for specific detection of *C. michiganensis* species. In both cases, the probe is located in the inner part of the amplicon, between the sense and anti-sense primers.

**S3 Fig.** Effect of 5’ AT-rich flap primers on ten-fold serially diluted sensitivity assays performed in endpoint PCR. **(A)** and **(B)** Multiplex reactions performed for detection of subspecies within the *C. michiganensis* and *C. m.* subsp. *nebraskensis* with/without 5’ AT-rich sequences, respectively; (**C)**. Single reactions targeting MSF gene of *C. m.* subsp. *nebraskensis*; **(D)** Single reactions targeting the sugar ABC transporter permease and ABC transporter ATP-binding genes for specific detection of *C. michiganensis* subspecies; **(E)** 1 µl host corn DNA was added in each reaction of ten-fold serially diluted sensitivity assay for simultaneous detection of *C. m.* subsp. *nebraskensis* and other C. michiganensis subspecies. All the experiments were conducted the same day using genomic DNA from *C. m.* subsp. *nebraskensis*. A molecular-weight size marker of 100 bp from BioLabs was used as a standard reference to determine the product size. All PCR products were electrophoresed in a 3% agarose gel.

