## Supplementary figures and images for "Synergetic effect of non-complementary 5’ AT-rich sequences on the development of a multiplex TaqMan real-time PCR for specific and robust detection of *Clavibacter michiganensis* and *C. michiganensis* subsp. *nebraskensis*"

### S1 Fig.

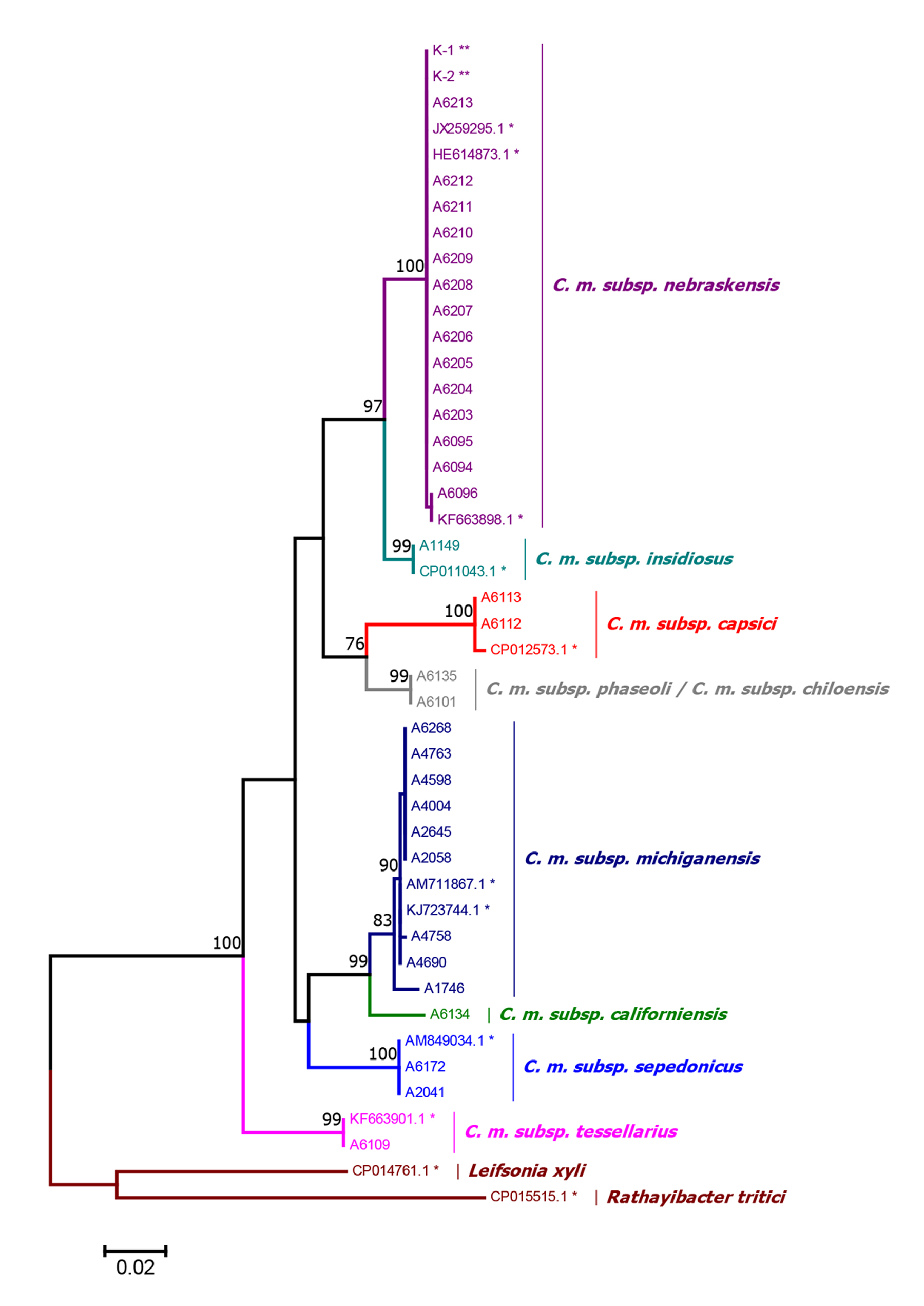

### S2 Fig.

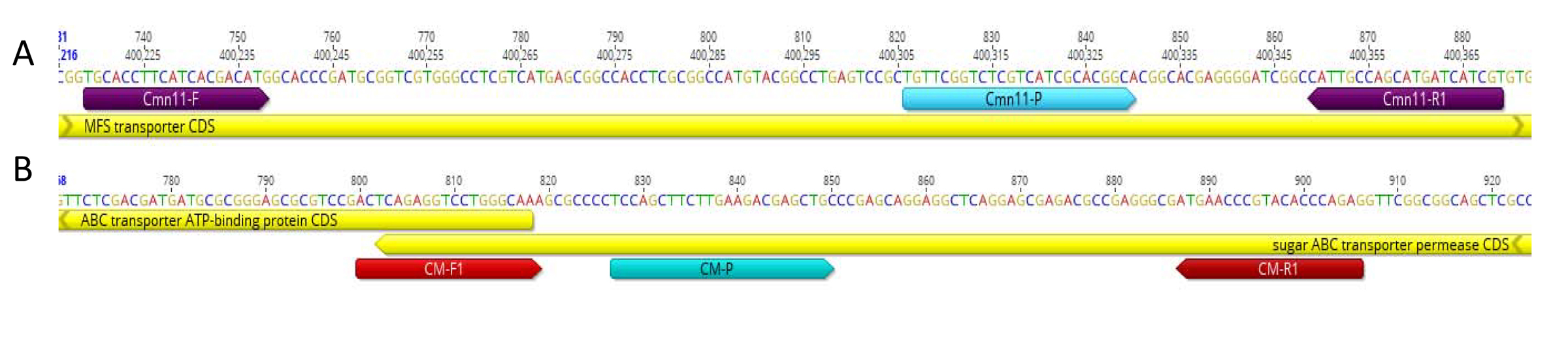

### S3 Fig.

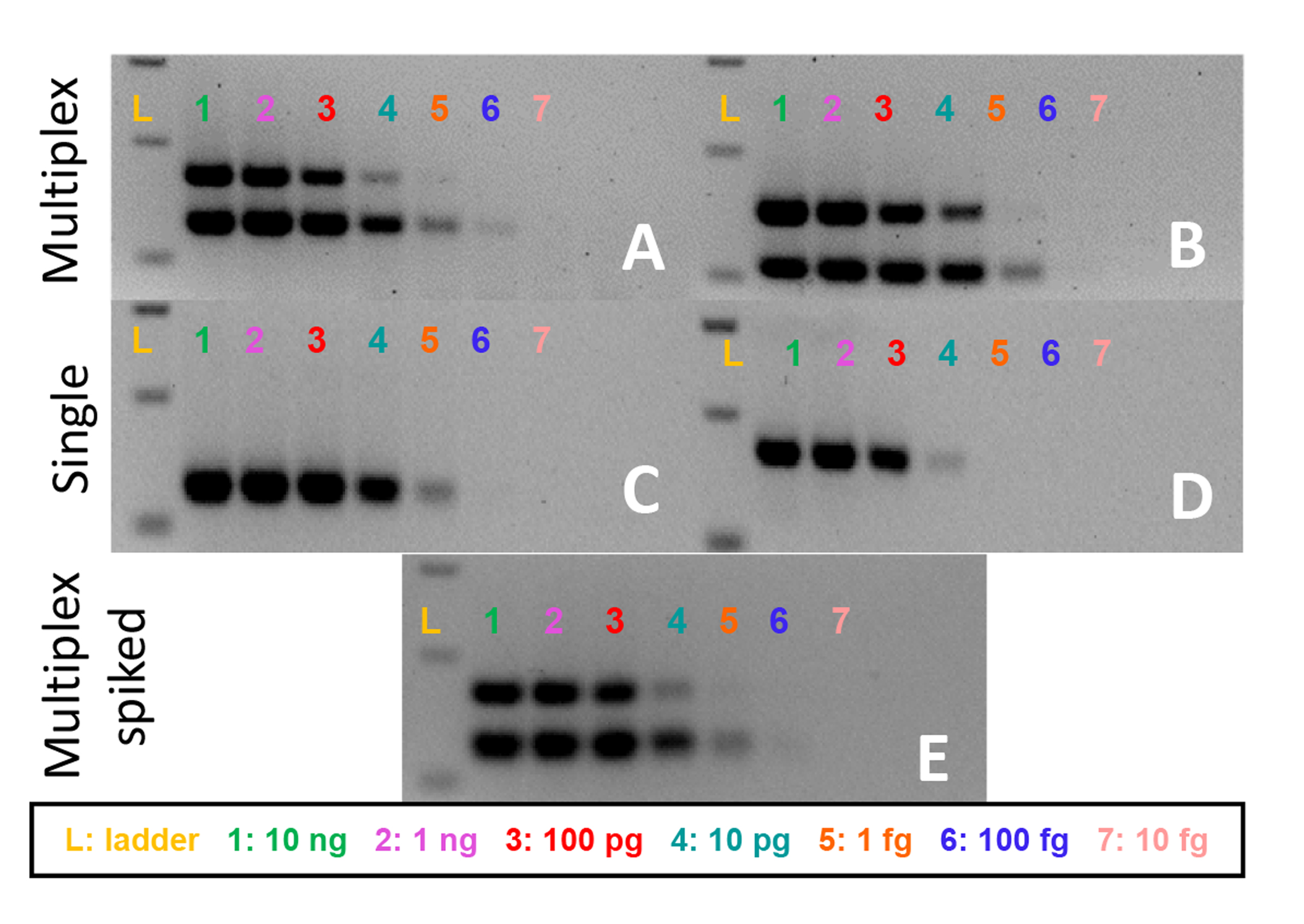
